# Endoplasmic reticulum-targeting but not translation is required for mRNA balancing in trypanosomes

**DOI:** 10.1101/2021.05.05.442555

**Authors:** Erick O Aroko, Majeed Bakari Soale, Christopher Batram, Nicola G Jones, Markus Engstler

**Affiliations:** Department of Cell and Developmental Biology, University of Würzburg, Würzburg, Germany

## Abstract

The cell surface of bloodstream form African trypanosomes is covered by a dense coat of immunogenic variant surface glycoproteins (VSGs). By continuously changing the expressed VSG antigen, the parasites can survive the host’s immune response. The *VSG* is highly expressed in *Trypanosoma brucei*, accounting for approximately 10 – 20% of total mRNA. Depletion of *VSG* mRNA is lethal, and a counterbalancing of the mRNA levels occurs when two or more *VSGs* are simultaneously expressed. How the VSG expression levels are regulated is unknown. Here, by using inducible and constitutive systems for ectopic VSG expression, we have discovered that (i) the endogenous *VSG* mRNA level is downregulated only when the ectopic *VSG* is targeted to the ER, (ii) VSG translation is dispensable and in fact, (iii) the regulation of *VSG* mRNA levels does not depend on a *VSG* open reading frame. We propose that feedback elicited at the ER regulates the *VSG* mRNA amounts to avoid overshooting the secretory pathway capacity. In this way, VSG expression is quantitatively and qualitatively fine-tuned. Balancing the overall number of ER-targeted mRNAs could well be a general mechanism in cell biology. The trypanosome system with just one dominant mRNA species provides a versatile model for studying this phenomenon.

## Introduction

African trypanosomes parasitize humans and a wide range of mammals, causing sleeping sickness and Nagana, respectively. These parasites have a digenetic life cycle, shuttling between the mammalian hosts and their insect vector — the tsetse fly. The mammalian stage bloodstream form (BSF) parasites are extracellular, residing in the blood, lymph, and the interstitial fluid of various tissues. Thus, African trypanosomes are under constant attack from the host’s immune system. Therefore, their cell surface has evolved as first and utmost important defense line. The cell surface of BSF *Trypanosoma brucei* is the best characterized compared to other African trypanosomes. This parasite is coated by a dense monolayer of an immunogenic glycosylphosphatidylinositol (GPI) anchored protein known as the variant surface glycoprotein (VSG). Other than providing an impenetrable protective shield to the parasites, the VSG coat is essential for antibody clearance and antigenic variation (Engstler et al., 2007; Mugnier et al., 2015; Pays et al., 2004). Together, these mechanisms ensure that the parasites stay ahead of the host’s immune responses.

Most proteins that traverse the eukaryotic secretory pathway, including VSGs, are flanked by endoplasmic reticulum (ER) import signal peptides at their N-terminal ends (Kreil, 1981; McConnell et al., 1981). Although a consensus sequence for naturally occurring ER import signals has not been defined, they contain some shared features. The ER import signals consist of a positively charged N-terminal region (n-region), a hydrophobic core (h-region), and a polar C-terminal (c-region) (Hegde & Bernstein, 2006; Kunze & Berger, 2015). Variations in the signal sequences affect different aspects of protein biogenesis including, targeting and translocation, cleavage, and ER exit (Hegde & Bernstein, 2006). The order of residues in the n-region is shown to influence the translocation of some substrates (Owji et al., 2018), and the hydrophobicity of the h-region is reported to be vital in determining whether a substrate is translocated by the signal receptor particle (SRP)-dependent or independent pathway in yeast (Ng et al., 1996). Further, the length and overall hydrophobicity of the h-region seem to be critical for translocation (Nilsson et al., 2015). *T. brucei* VSG ER import signal peptides are approximately 15 – 30 residues long and share the signal peptide features described in yeast and mammals (Boothroyd et al., 1981; Cross, 1984; McConnell et al., 1981).

Transcription of virtually all eukaryotic genes is monocistronic. However, most *T. brucei* protein-coding genes are arranged in polycistronic arrays transcribed by RNA polymerase II (Johnson et al., 1987; Siegel et al., 2011). The long polycistronic transcripts are processed by trans-splicing a 39-nucleotide spliced leader at the 5′ end, and 3′ polyadenylation of individual genes to form mature mRNAs (Michaeli, 2011). Due to this polycistronic arrangement of genes and a lack of transcriptional initiation control of individual genes, regulation of RNA levels in *T. brucei* is predominantly post-transcriptional (Clayton, 2002). RNA-binding proteins (RBPs) and sequence elements within the 3′UTRs of genes are crucial for this post-transcriptional modulation of *T. brucei* gene expression (Clayton, 2019; Kramer, 2012). Transcription of the major surface proteins in *T. brucei* is unusual in that it is driven by RNA polymerase I. In the BSF, a single *VSG* gene located at the subtelomeric locus of one of ∼15 polycistronic VSG expression units known as the expression site (ES), is monoallelically expressed to ensure that only a single VSG is produced at a time from the vast repertoire of 2000 genes and pseudogenes (Cross et al., 2014; Hertz-Fowler et al., 2008). The active ES localizes to the extranucleolar expression site body (ESB) in the nucleus (Budzak et al., 2019; Navarro & Gull, 2001).

The expression of VSG is tightly regulated to ensure an intact coat is maintained both *in vivo* and *in vitro*. Down-regulation of VSG expression via RNAi or by blocking translation using morpholino oligonucleotides causes a precise pre-cytokinesis cell cycle arrest and subsequent cell death (Sheader et al., 2005(Barnwell et al., 2010; Ooi et al., 2018). A study by (Muñoz-Jordán et al., 1996) showed *T. brucei* could be engineered to constitutively express two VSGs in equal amounts by integrating a second *VSG* gene just upstream of the active ES-resident *VSG*. Integration of a second *VSG* gene downstream of the active ES promoter — approximately 60 kb upstream of the native *VSG* — also supported stable expression of two VSGs with roughly equal amounts of both mRNA and protein, that did not exceed the wild-type levels. In addition, when transcripts of one of the *VSGs* were depleted using RNAi, there was a compensatory effect by which the transcripts of the other *VSG* were upregulated to wild type levels (Smith et al., 2009). A similar apparent upregulation of the ectopic *VSG* expression was observed when the native ES-resident *VSG* was deleted by replacement with a blasticidin resistance gene (Ridewood et al., 2017). Expression site residence is not required for *VSG* mRNA regulation. Inducible overexpression of an ectopic *VSG121* from the ribosomal spacer region resulted in a fast and efficient exchange of the *VSG* mRNA population and protein, with similar kinetics in both monomorphic and pleomorphic trypanosomes (Batram et al., 2014; Zimmermann et al., 2017). In the monomorphic cell line, the ectopic *VSG* mRNA is rapidly upregulated to 80% of the wild type levels within 2 h of inducing expression and this level of expression is maintained for 24 h. Concomitantly, the endogenous *VSG* mRNA is downregulated and reaches ∼25% of wild type amounts 6 h after inducing expression (Batram et al., 2014). Taken together, these data suggested the existence of a robust mechanism, possibly involving regulatory steps at both transcript and protein level, to ensure a constant level of VSG expression required for maintaining a flawless VSG coat.

One candidate signal that could be involved in *VSG* mRNA control is a conserved 16-mer sequence present in all *T. brucei* VSG 3′UTRs — essential for high expression and stability of *VSG* mRNA (Berberof et al., 1995). This sequence could be bound by a putative limiting RBP and somehow counted (Ridewood et al., 2017). Recently, it has been proposed that this limiting factor could be the 16-mer motif-dependent inclusion of *N*^*6*^-methyladenosine in the *VSG* poly(A) tails (Viegas et al., 2020). In another recent study an mRNA binding protein CFB2 was found to interact with the 16-mer motif, where it mediates VSG mRNA stability by recruitment of a stabilizing complex. In addition, it was proposed that regulation of CFB2 levels might in turn limit VSG synthesis to avoid excess production (do Nascimento et al., 2020). However, by introducing premature termination codons (PTCs) in the ectopic *VSG* gene open reading frame (ORF), Maudlin et al. reported massively increased total *VSG* mRNA levels thus questioning whether a direct counting mechanism for *VSG* mRNA exists (Maudlin et al., 2021).

Here, we show that the endogenous *VSG* mRNA regulation response is elicited only when the ectopic VSG is correctly targeted to the ER. Additionally, we found that there is nothing special about the VSG transcript in terms of autoregulation of its abundance, and that high level expression of any ER targeted mRNA is sufficient to elicit this VSG regulation response. Thus, the counterbalancing of *VSG* mRNA levels appears to be a strategy for regulating the secretory pathway’s cargo and maintaining ER homeostasis. As the regulation is independent of the open reading frame, it could well facilitate the rapid exchange of surface coats during antigenic variation and developmental progression.

## Results

### Overexpression of a *T. vivax* VSG gene causes the downregulation of endogenous *T. brucei VSG* mRNA

The amount of *VSG* mRNA in *Trypanosoma brucei* appears to be tightly regulated, potentially due to the presence of the 16-mer motif in the 3′UTR, which is the common denominator of all VSGs in this trypanosome species. Antigenic variation of VSG, however, is not restricted to the *brucei*-group of trypanosomes but is also present in the related species *T. congolense* and *T. vivax*. However, *T. congolense* and *T. vivax* VSGs lack the conserved 16-mer element. Furthermore, these VSGs are smaller than the *T. brucei* VSGs and the *T. vivax* VSG coat may not be as dense as the *T. brucei* coat (Autheman et al., 2021; Gardiner et al., 1987). Therefore, it is unclear if and how *T. congolense* and *T. vivax* use *VSG* mRNA balancing. To address this, we expressed *T. vivax* VSG genes in *T. brucei* to test whether *T. brucei* specific sequence features are required for the counterbalancing of *VSG* mRNA amounts. For this purpose, we selected two well documented *T. vivax* VSGs — *ILDat1*.*2*: *TvY486_0008160* and *ILDat2*.*1*: *Z48228*.*1* (Barry & Gathuo, 1984; Chamond et al., 2010; Gardiner et al., 1987; Gardiner et al., 1996; Greif et al., 2013) — for inducible overexpression in *T. brucei*. This expression strategy was used because it enables monitoring of early events after induction of VSG expression, and it is not obscured by adaptive responses to constitutive ectopic VSG expression, which can occur during selection of transgenic parasites. First, a construct for the expression of ILDat2.1 VSG was generated. As the published *ILDat2*.*1* sequence lacks a start codon (Gardiner et al., 1996), we added a start codon to the *ILDat2*.*1* VSG open reading frame. Given that the conserved elements in the VSG 3′UTRs are unique to *T. brucei* and essential for high levels of expression (Berberof et al., 1995; Ridewood et al., 2017), a *T. brucei* VSG 3′UTR sequence was fused to the *ILDat2*.*1 VSG* ORF, which was then integrated into the transcriptionally silent rDNA spacer region using the pLew82v4 vector (24 009; Addgene plasmid). Expression of the ectopic VSG genes was driven by an ectopic tetracycline inducible T7 RNA polymerase promoter (Figure 1A) (Batram et al., 2014).

**Figure 1:**
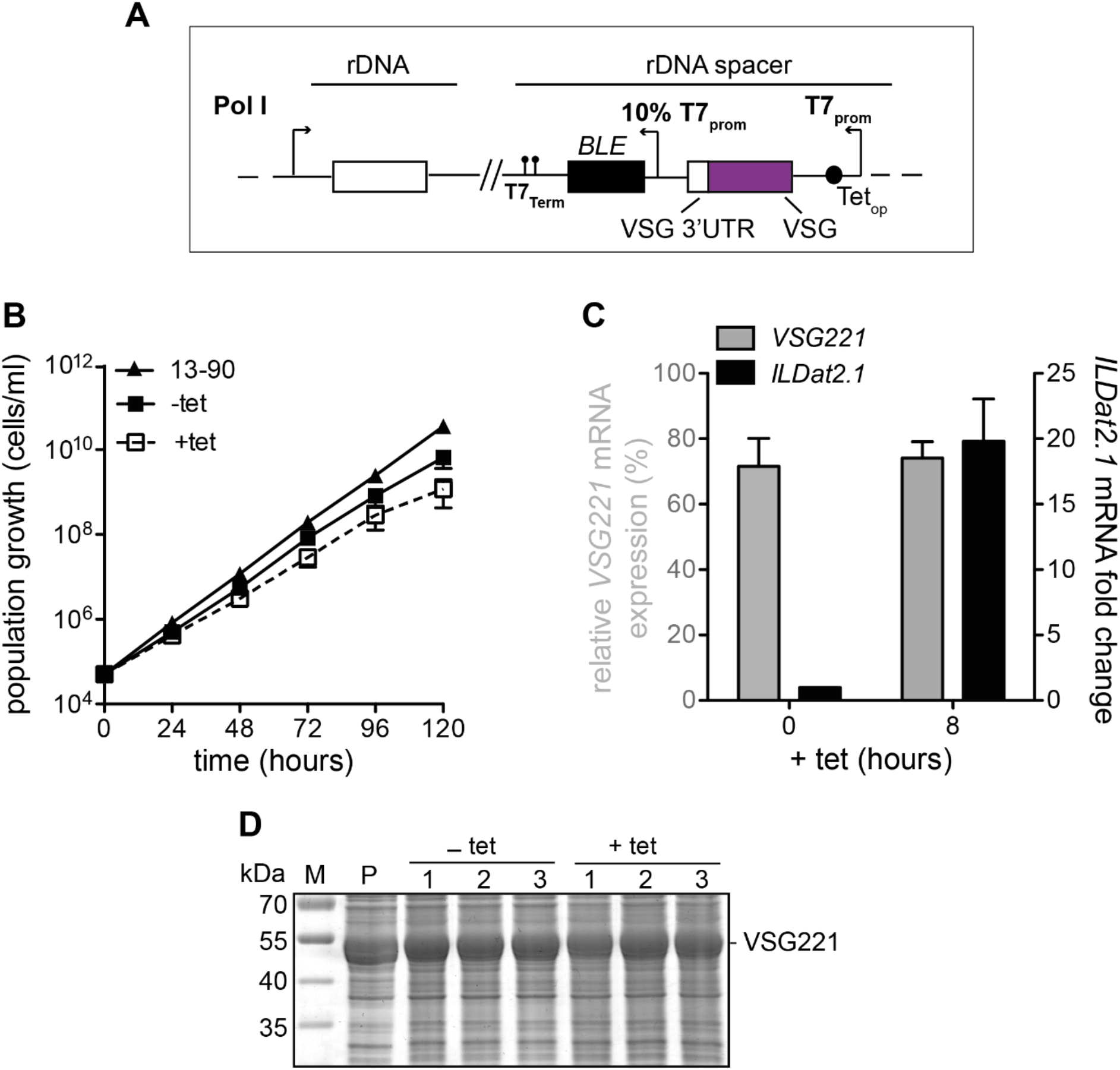
Endogenous VSG mRNA regulation response is not elicited upon Induction of ILDat2.1 expression. (A) Schematic of the ectopic VSG overexpression strategy. The *ILDat2*.*1* construct was integrated into the transcriptionally silent rDNA spacer and expression of the VSG was driven by the T7 promoter in the presence of tetracycline. (B) Cumulative growth profiles of 221^ES^.ILDat2.1^tet^ cells. Three independent clones were analyzed for 5 days in the presence (+tet) or absence (-tet) of tetracycline. The parental 13-90 cells (P) served as a growth control. (C) Quantification of *VSG* mRNA levels in the presence (8 h) or absence (0 h) of tetracycline. Expression of *VSG221* (left y-axis) is expressed as a percentage relative to the parental 13-90 cells *VSG221* expression while the ectopic *ILDat2*.*1* value (right y-axis) is relative to the non-induced levels. The values are given as mean ± SD (Standard deviation) of three independent clones. The mRNA was normalized to *β-tubulin*. (D) Analysis of ILDat1.2 protein expression by SDS-PAGE after 24 h in the presence (+tet) or absence (-tet) of tetracycline. The parental 13-90 cells (P) served as the loading control and three independent clones (1 – 3) were analyzed.

Inducing the expression of ILDat2.1 VSG in the 221^ES^.ILDat2.1^tet^ cell line caused a minor reduction in proliferation (Figure 1B). We tested whether high levels of *ILDat2*.*1 VSG* mRNA were transcribed and if so, whether an endogenous *VSG* mRNA regulation response was triggered upon induction of expression. It is known that this expression system is leaky, causing 10 – 20% expression of ectopic VSGs relative to the wild type VSG levels before inducing expression. Quantitative dot blots carried out 8 h after induction showed approximately 20-fold increase of the *ILDat2*.*1 VSG* mRNA. This increase was higher than the approximate 10-fold change recorded when wild type *T. brucei* VSGs are expressed in this way. The endogenous *VSG221* mRNA level was, however, not downregulated (Figure 1C). This is in contrast to the regulation that occurs when *T. brucei* VSGs are overexpressed. Additionally, the ILDat2.1 VSG protein appeared not to be expressed at all (Figure 1D). As the endogenous *VSG221* mRNA was not affected, and thus VSG221 protein levels remained high, the transgenic trypanosomes grew normally. High level expression of a *T. vivax VSG* mRNA had no effect on the ES-resident VSG, which could mean that there are, in fact, special features in the *T. brucei* VSG open reading frame that are required for mRNA balancing.

To test this possibility, a second *T. vivax* VSG (ILDat1.2) containing a *T. brucei* VSG 3′UTR was used. Induction of expression of the ILDat1.2 VSG in the 221^ES^.ILDat1.2^tet^ cell line, led to significant slowing in growth within 24 h and complete stalling thereafter (Figure 2A). Quantitative dot blots carried out 8 h after inducing expression showed that a high amount of *ILDat1*.*2 VSG* mRNA was present, with a concomitant reduction of the endogenous *VSG221* mRNA to 20% of the wild type levels (Figure 2B). This phenotype contrasted the above results obtained with VSG ILDat2.1, which was difficult to interpret. Analysis of ILDat1.2 expression showed that the protein was produced in only low amounts (Figure 2C), differing from what happens when *T. brucei* VSGs are overexpressed. This means that downregulation of the endogenous *VSG* mRNA, in addition to the insufficient production of the ectopic ILDat1.2, resulted in a shortage of VSG protein, slowed parasite growth and eventual cell death. In search of the reason for the poor production of ILDat1.2 VSG, we surmised that the VSG GPI-anchoring signal might not be adapted to function in *T. brucei*. Supporting this assumption, the phenotype was rescued by replacement of the native ILDat1.2 GPI anchor signal with that of a *T. brucei* VSG. This ILDat1.2 VSG transgene was highly expressed on both mRNA and protein levels. The abundant ILDat1.2 VSG protein (Figure 2D) rescued the growth phenotype (Supplementary Figure 3).

**Figure 2.**
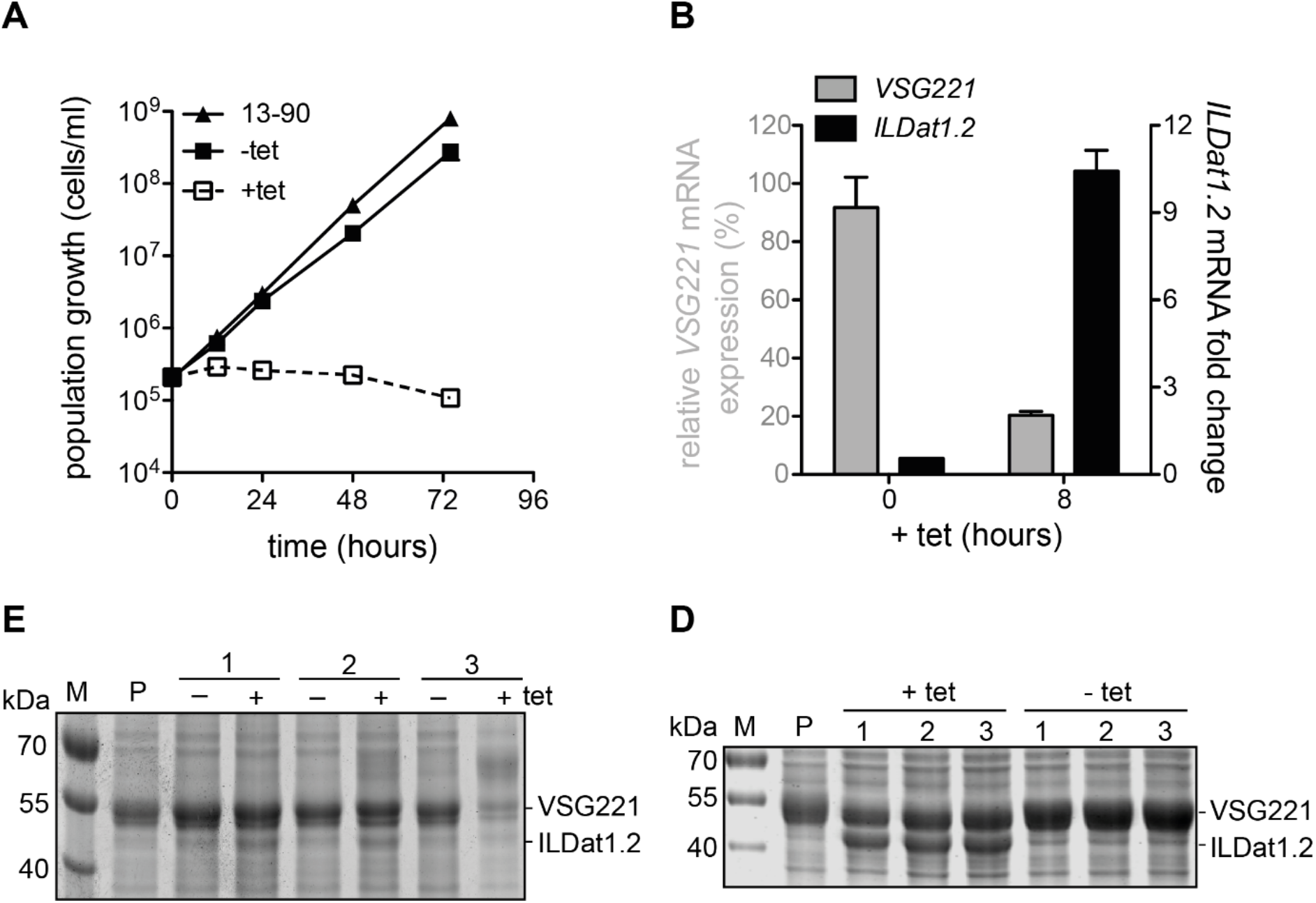
The endogenous regulation response is elicited upon induction ILDat1.2 VSG expression. (A) Cumulative growth of the 221^ES^.ILDat1.2^tet^ cell line in the presence (+tet) or absence (-tet) of tetracycline and the parental 13-90 cells (P) served as a growth control. (B) Quantification of mRNA levels of the ectopic *ILDat1*.*2* expressed from the rDNA spacer and the endogenous *VSG221* using dot blots. The cells were induced to express the ectopic *VSG* for 8 h. Expression of *VSG221* (left y-axis) is given as a percentage relative to the parental 13-90 cells *VSG221* expression while the ectopic *ILDat1*.*2* value (right y-axis) is relative to the non-induced levels. *VSG* transcript levels were normalized to *β-tubulin* and the results given as means of three independent clones. The error bars indicate the SD. (C) SDS-PAGE analysis of wild type ILDat1.2 and (D) ILDat1.2 with a *T. brucei* GPI signal protein expression after 24 h in the presence (+tet) or absence (-tet) of tetracycline. The parental 13-90 cells (P) served as the loading control and three independent clones were analyzed.

The above results clearly demonstrate that, in principle, *T. vivax VSG* mRNA that is coupled to a *T. brucei* 3′UTR can elicit *VSG* mRNA balancing in *T. brucei*. Nevertheless, the *VSG* mRNA *trans*-regulation mechanism was only operational in the ILDat1.2 overexpression cell line but not in the ILDat2.1 overexpression cell line, leaving us with the question why.

### ER-targeting but not translation is essential for the *trans*-regulation of *VSG* mRNA levels

To solve this puzzle, we initially replaced the native GPI anchor signal of ILDat2.1 with that of a *T. brucei* VSG. This, however, neither lead to regulation of the endogenous *VSG* mRNA nor high-level ectopic VSG protein expression, despite the production of high levels of *ILDat2*.*1* mRNA. The start codon of *ILDat2*.*1 VSG* was missing in the published sequence data (Gardiner et al., 1996) and signal peptide prediction software revealed *ILDat2*.*1* to have a low probability score for the presence of an ER import signal. ER import signals are reported to influence different aspects of the biogenesis of secretory proteins. Features found within the signals, including hydrophobicity, differentially influence processing of nascent polypeptides (Hegde & Bernstein, 2006). We compared the hydrophobicities of the ER signal peptides of *T. vivax* VSGs with those of *T. brucei* VSGs and non-VSG surface proteins. The hydropathy characteristics were calculated as the grand average hydropathy (GRAVY) score of all the amino acids in the signal peptide (Kyte & Doolittle, 1982). The reported ILDat2.1 ER import signal had a negative GRAVY score, indicating that it was hydrophilic, whereas the other analyzed ER import signals had positive scores. Further, the hydrophobic core of the ILDat2.1 ER signal peptide could not be predicted using Phobius (Käll et al., 2007) as the signal was not recognized (Supplementary Table 1). Additionally, the published ILDat2.1 ER import signal (Gardiner et al., 1996) does not appear to function in *T. brucei* and, based on ER import signal characteristics, may not actually be a functional ER import signal at all. These data support our hypothesis that for the endogenous *VSG* mRNA to be regulated, the ectopic *VSG* transcript must be targeted to the ER. Therefore, we surmised that the ILDat2.1 VSG ER import signal may lead to a mistargeting of the *ILDat2*.*1* mRNA.

We performed two experiments to test this possibility: (i) we tried to express VSG121 with the potentially defective ILDat2.1 ER import signal (Figure 3A, upper panel), and (ii) we tried to rescue expression of ILDat2.1 by replacing the ILDat2.1 ER import signal with that of *T. brucei* VSG121 (Figure 3A, lower panel).

**Figure 3.**
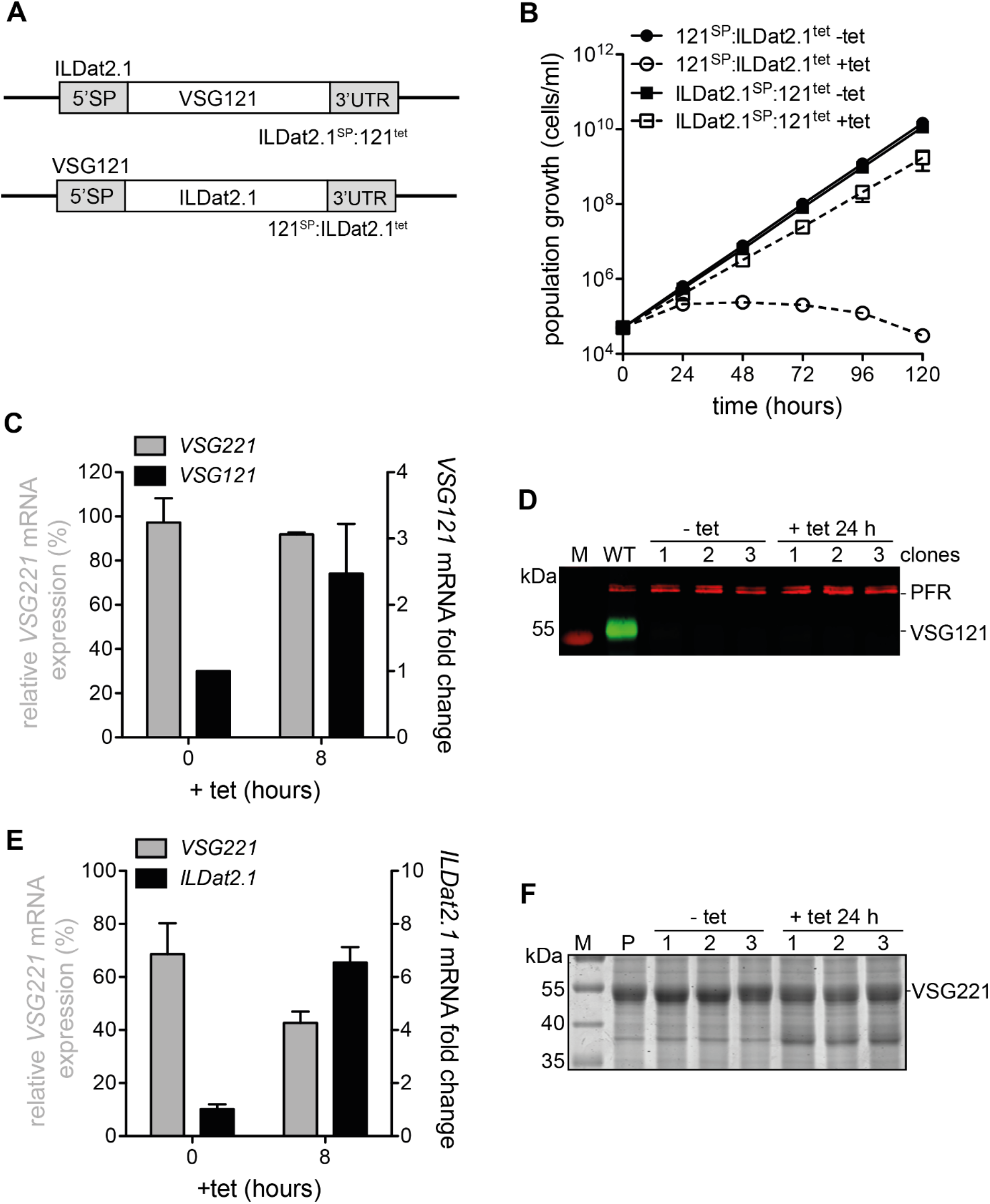
The endogenous *VSG* mRNA is regulated only when the ectopic VSG has a functional ER-import signal. (A) Schematic of the constructs used to generate the 121^SP^:ILDat2.1^tet^ (upper panel) and ILDat2.1^SP^:121^tet^ (lower panel) cell lines. (B) Cumulative growth curves of the 121^SP^:ILDat2.1^tet^ and ILDat2.1^SP^:121^tet^ cell lines. Three independent clones were analyzed for 5 days in the presence (+tet) or absence (-tet) of tetracycline. (C) Quantification of *VSG* mRNA levels before (0 h) and after (8 h) inducing expression of the ectopic *VSG* in the ILDat2.1^SP^:121^tet^ cells. Expression of *VSG221* (left y-axis) is expressed as a percentage, relative to the parental (P) 13-90 cells *VSG221* expression while the *VSG121* value (right y-axis) is relative to the non-induced levels. The mRNA was normalized to *β-tubulin*. Two independent clones were analyzed, and the values are presented as means ± SD. (D) Western blot analysis of VSG121 protein expression in ILDat2.1^SP^:121^tet^ cells before (-tet) and 24 h after (+tet) inducing expression using anti-VSG121 antibodies. Wild type (WT) *T. brucei* cells expressing VSG121 served as the positive control (green) and PFR protein (red) served as the loading control. (E) Analysis of *VSG* mRNA expression in three clones of the 121^SP^:ILDat2.1^tet^ cells. The endogenous *VSG221* and ectopic *ILDat2*.*1* are quantified and presented as in C. (F) SDS-PAGE analysis of ILDat2.1 protein expression in the 121^SP^:ILDat2.1^tet^ cells before (-tet) and 24 h after (+tet) addition of tetracycline. The 13-90 parental cells (P) served as the loading control.

Induced expression of high amounts of the ILDat2.1^SP^:121 fusion mRNA did not affect parasite growth (Figure 3B). Likewise, the endogenous *VSG221* mRNA levels in the ILDat2.1^SP^:121^tet^ cell line were not downregulated (Figure 3C). No *VSG* mRNA balancing was observed. Protein expression analysis revealed that the VSG reporter was not expressed (Figure 3D). Since endogenous VSG mRNA and protein were expressed abundantly, the cells grew well. This result confirmed our assumption that the ILDat2.1 ER signal was defective, which was further supported by the second experiment. Inducing the expression of ILDat2.1 VSG fused to the VSG121 ER import signal led to markedly impaired cell growth within 24 h, followed by cell death. Analysis of *VSG* transcripts in the 121^SP^:ILDat2.1^tet^ cell line showed that expression of the VSG chimera caused a simultaneous reduction of the endogenous *VSG221* mRNA to ∼40% of the wild type levels within 8 h (Figure 3E). Interestingly, despite the transcription of the reporter *VSG* mRNA, only a faint protein band of between 35 – 40 kDa was observed, indicating that the recombinant *T. vivax* VSG protein was poorly expressed (Figure 3F). The marked decline in endogenous *VSG* mRNA resulted in depletion of wild type VSG protein which eventually resulted in cell death (Figure 3B). While we still have no explanation for the failure of the abundant 121^SP^:ILDat2.1 mRNA to yield high protein levels, the experiments fortuitously suggested a new hypothesis on the prerequisites for *VSG* mRNA balancing: (i) A *VSG* mRNA from another trypanosome species (*T. vivax* ILDat1.2) is functional in mRNA balancing. (ii) The *VSG* mRNA has to be successfully targeted to the ER. (iii) The *VSG* mRNA does not have to be translated to trigger mRNA balancing. These surprising insights led us to raise two further questions. First, is a bona fide VSG ER signal sequence required and, second, is the presence of a bona fide *VSG* mRNA (coming from any VSG coated trypanosome species) required for the balancing. In other words, is the signal hidden in the VSG ORF?

### Non-VSG ER import signals can target VSGs to the ER

We first tested the possibility that the VSG ER signal itself has unique features. We selected the ER import signals of abundant GPI-anchored surface proteins of different insect stages of *T. brucei* (including the epimastigote-specific *brucei* alanine-rich protein (BARP), and the procyclic EP1 protein) as well as the non-abundant procyclic factor H receptor (FHR). These surface proteins, like the VSGs, traverse the secretory pathway. Thus, we generated *VSG121* transgenes with BARP (C9ZZN9, UniProt), EP1 (Q389V1, UniProt) and FHR (Q57Z47, UniProt) ER import signals. This time we decided to use constitutive expression from the VSG 221 expression site, as mRNA balancing in VSG double expressors is not always 50:50 but may vary depending on subtle differences between VSGs. We have shown that integrating *VSG121* downstream of the *VSG221* results in a 70:30 expression of *VSG221* and *VSG121* mRNA, respectively, relative to the wild type amounts (Supplementary Figure 2). This demonstrates that balancing of *VSG* mRNAs is an intrinsic regulatory mechanism that is not just modulated by transcription rates. Thus, the “quality” of the ER targeting could be reflected in the mRNA ratios between the native VSG221 and the VSG121 chimera. We integrated the recombinant VSG genes just downstream of the active ES resident *VSG221* to generate the cell lines 221^ES^.BARP^SP^:121, 221^ES^.EP1^SP^:121 and 221^ES^.FHR^SP^:121 (Figure 4A).

**Figure 4.**
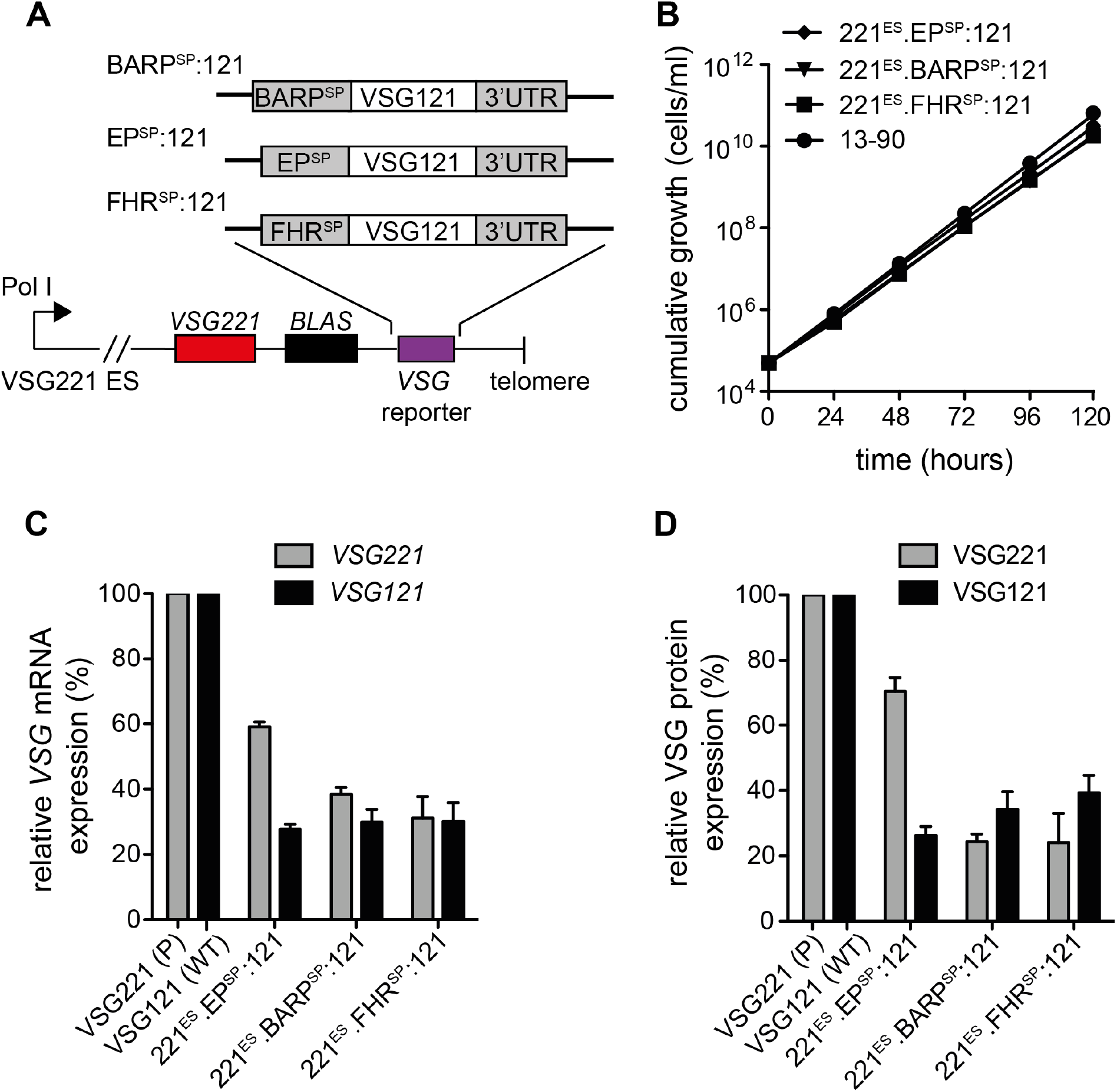
Non-VSG ER import signals can target VSGs to the ER. (A) Illustration of the constructs and expression strategy used to generate the 221^ES^.BARP^SP^:121, 221^ES^.EP^SP^:121 and 221^ES^.FHR^SP^:121 cell lines. (B) Analysis of the growth curves showed a slight reduction of growth in all of the generated cell lines as compared to the parental 13-90 cells. The 221^ES^.BARP^SP^:121 and 221^ES^.FHR^SP^:121 cell lines appear as a single curve in the graph. (C) Quantification of *VSG* mRNA and (D) protein levels using dot blots. *VSG221* mRNA expression levels are relative to the parental (P) 13-90 *VSG221* expression levels while *VSG121* expression is relative to the Wild type (WT) *VSG121* expression levels. The mRNA was normalized to *β-tubulin* whereas the VSG protein was normalized to the PFR protein. The values are presented as means ± SD of three independent clones.

The three experiments yielded clonal trypanosome cell lines that did not display a growth phenotype (Figure 4B). At the transcript level, a *trans*-regulation of the *VSG* mRNA was elicited in all three clones. In the 221^ES^.BARP^SP^:121 cell line, the endogenous *VSG221* and the ectopic *VSG121* were expressed at 40% and 30% of the wild type levels, respectively, whereas both *VSGs* were approximately 30% of the wild type levels in the 221^ES^.FHR^SP^:121^tet^ cell line. In the 221^ES^.EP1^SP^:121 cell line, the endogenous *VSG221* and the ectopic *VSG121* were expressed at 60% and 30% of the wild type levels, respectively (Figure 4C). Although the *trans*-regulation mechanism was operational in the 221^ES^.BARP^SP^:121 and 221^ES^.FHR^SP^:121 cells, the total *VSG* mRNA was ∼70% of the wild type levels and the ectopic VSG121 protein appeared to be expressed more than the endogenous VSG221 (Figure 4D). Together, these findings indicated that the ER import signals of non-VSG surface proteins can target VSGs to the ER. However, it appears the ER signals varyingly influence the regulation of *VSG* mRNA levels.

### The *VSG* open reading frame is not required for trans-regulation of *VSG* expression

The above experiments suggested that balancing of *VSG* mRNA requires targeting the transcripts to the ER. Though this balancing is independent of VSG-specific ER signals, non-VSG ER signals appear to influence it differently. The finding that divergent VSGs from *T. vivax* can elicit *VSG* mRNA balancing can be interpreted in two ways: First, *T. vivax* VSGs can functionally complement *T. brucei* VSGs, or, second, mRNA balancing is independent of the ORF and just requires abundant mRNA targeted to the ER.

To test these possibilities, we used the overexpression strategy to express high levels of GFP reporters in cells with an active *VSG221* expression site. The first reporter consisted of the *GFP* ORF and the *VSG121* 3′UTR. This construct was integrated into the rDNA spacer to generate the 221^ES^.GFP^tet^ cell line that inducibly expresses GFP in the cytosol. We also generated a second reporter for expression of ER-targeted GFP. In addition to the *VSG* 3’UTR sequence, a procyclin (EP1) ER import signal was coupled to the 5′ end of the *GFP* ORF and integrated into the transcriptionally silent rDNA spacer, yielding the 221^ES^.EP^SP^:GFP^tet^ cell line that inducibly expresses an ER-targeted GFP reporter. If neither the *VSG* ORF nor a specific ER import sequence were required for VSG balancing, the expression of high levels of ER-targeted *GFP* should lead to a modulation of *VSG* mRNA.

After inducing expression, a strong GFP signal was detected in the cytoplasm of the cell line expressing the GFP reporter lacking the ER signal by fluorescence microscopy (Figure 5A). On the other hand, a patchy expression of GFP was observed in a compartment consistent with the ER after inducing the expression of ER targeted GFP (Figure 5B). A slight reduction in growth was recorded within the first 24 h of cytosolic reporter overexpression, after which the cell numbers began to decline (Figure 5C), possibly due to the toxicity of high levels of GFP (Liu et al., 1999). In the cell line expressing the ER-targeted reporter, a rapid stalling in parasite growth was observed after inducing expression, followed by a decline in cell numbers within 6 h (Figure 5D). As we were specifically interested in very early effects after induction, this later growth defect did not affect the interpretation of the data. On the transcript level, quantitative dot blots showed an immediate increase in the *GFP* mRNA in the 221^ES^.GFP^tet^ cell line, with peak *GFP* expression recorded between 2 – 6 h after induction, followed by a decline. There was no impact on the *VSG* mRNA levels within the first 6 h of inducing cytosolic GFP overexpression. However, after 8 h, there was a reduction of both the *VSG* and *GFP* mRNAs. The *VSG* transcript reverted to wild type levels after 24 h (Figure 5E). Interestingly, expression of the *GFP* transgene containing an ER signal caused a rapid simultaneous reduction of the *VSG* mRNA amount. The endogenous *VSG221* mRNA decreased to 20% of the wild type level within 2 h of expression (Figure 5F). The results of these experiments suggested that, in fact, the regulation of endogenous *VSG* neither required the ectopic *VSG* open reading frame nor the VSG ER signal.

**Figure 5.**
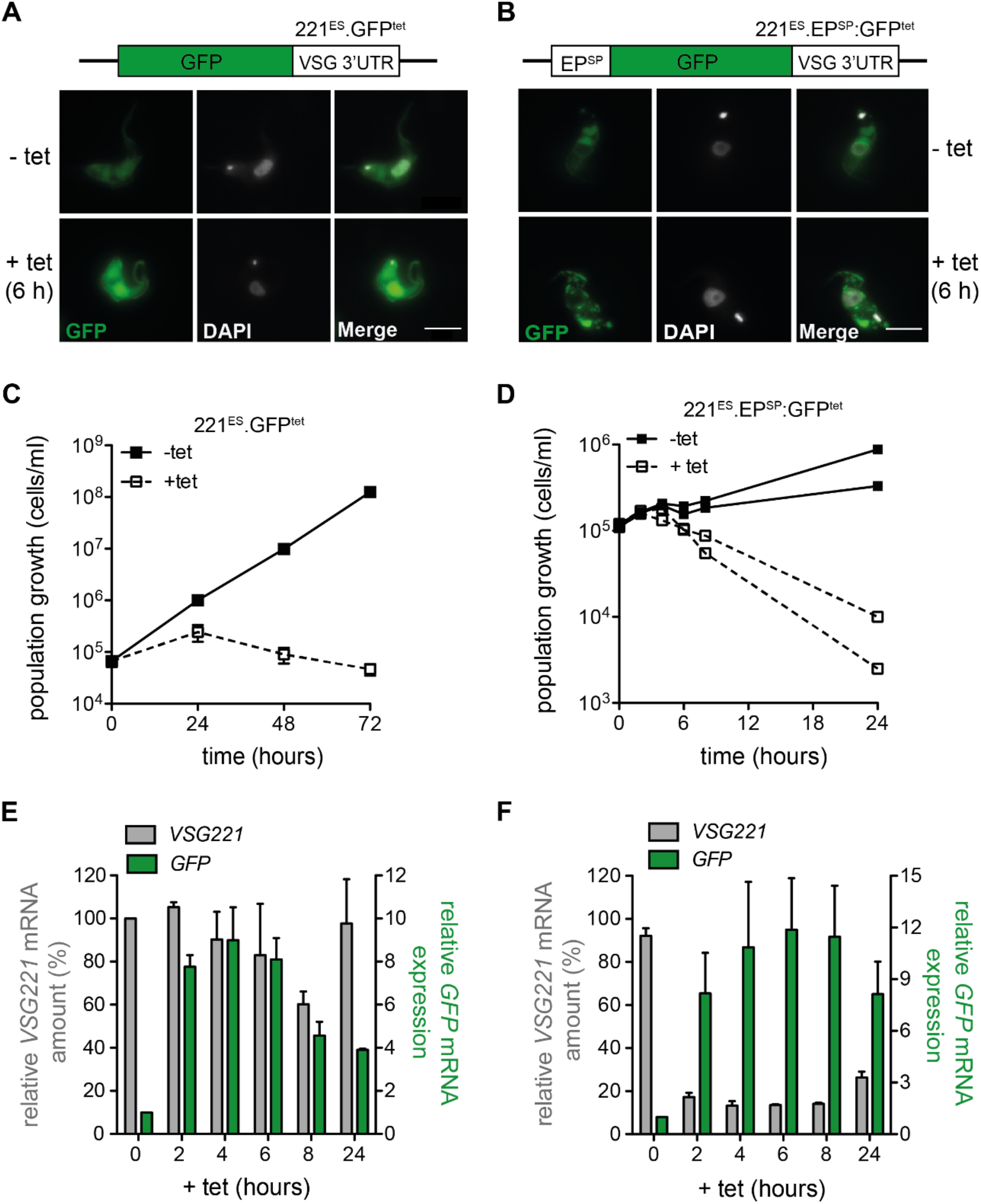
High-level expression of ER-targeted *GFP* reporter causes downregulation of *VSG* mRNA amounts. (A) Schematic of the GFP reporter constructs and fluorescence microscopy images of fixed cells showing the expression of GFP before (-tet) and 6 h post-induction of expression with 1 µg/ml tetracycline (+tet) in the 221^ES^.GFP^tet^ cells expressing cytosolic GFP, and (B) 221^ES^.EP^SP^:GFP^tet^ cell line expressing GFP in the ER. GFP fluorescence is shown in green while parasite nuclei and kinetoplasts were stained with 4,6-diamidino-2-phenylindole (DAPI, white). Scale bar: 5 µm. (C) Cumulative growth after induction of cytosolic *GFP* reporter expression. Three independent clones were analyzed for 72 h in the presence (+tet) or absence (-tet) of tetracycline (D) Cumulative growth of the cells expressing ER localized GFP reporter. Two independent clones were analyzed for 24 h. (E) Quantification of *VSG* (left y-axis) and *GFP* (right y-axis) mRNA levels during the course of tetracycline induced cytosolic and (F) ER targeted *GFP* reporter overexpression. Total RNA samples were dot-blotted and hybridized with fluorescently labelled probes specific for *VSG221, GFP* and *β-tubulin*. The data was quantified by normalization to *β-tubulin. VSG* mRNA expression is presented as percentage means ± SD for two independent clones normalized to the parental 13-90 cells expressing *VSG221* whereas *GFP* expression values are relative to the non-induced cells, -Tet = 1.

In conclusion we have shown that trypanosomes have evolved a fine-tuned and flexible mechanism to balance the number of ER-targeted mRNAs. This mechanism is independent of the nature of ER signal sequences and open reading frames and does not require translation. Thus, the trypanosome’s version of mRNA buffering supports the high phenotypic plasticity of the parasitic lifestyle.

## Discussion

Antigenic variation is a powerful strategy employed by several pathogens to evade elimination by host immune responses. Monoallelic expression of surface antigens by parasitic pathogens, such as trypanosomes and *Plasmodium falciparum*, ensure that only a single antigen is expressed at a time from a vast repertoire of silent genes (Borst, 2002; Voss et al., 2006). In trypanosomes, variation of the VSG antigens involves transcriptional switching or recombination events that introduce a new antigen into the active VSG expression site (Horn, 2014).

We have simulated an *in situ* VSG switch by inducible and high-level expression of a second VSG from the ribosomal spacer (Batram et al., 2014). Whatever VSG gene we expressed ectopically, we consistently found an almost instantaneous reaction from the expression site: the mRNA of the ES-resident VSG declined rapidly to low levels. We have further shown when overexpressing VSG121 in a VSG221 background that the parasites, in a next step, can attenuate the expression site in a DOT1b-dependent manner (Batram et al., 2014). This, however, is a stochastic process. Using a pleomorphic trypanosome strain, we found that in a clonal population, ES-silencing occurred in some 50% of all cells, while all cells silenced the VSG. This means that ES silencing is independent from VSG silencing (Zimmermann et al., 2017). Furthermore, not all ectopically expressed VSG genes were affecting the ES-resident *VSG* mRNA with equal efficiency, and not all ESs were equally responsive. Thus, trypanosomes exhibit a remarkable degree of phenotypic plasticity and high cell-to-cell variability. This makes perfect sense for a parasite that, during its life cycle, encounters extremely different microenvironments, and that does not regulate gene expression by transcriptional control as most other eukaryotes do. Instead, posttranscriptional control is paramount (Clayton, 2002).

In all our experiments, one observation was consistently made: the inducible expression of a second VSG only led to a very short period of overshooting total *VSG* mRNA, which then rapidly leveled to approximately wild type amounts (Batram et al., 2014; Zimmermann et al., 2017). We can observe the simultaneous increase of ectopic *VSG* mRNA and decline of ES-resident *VSG* mRNA. We have not only documented this by overexpressing VSGs, but also by generating so-called double expressor cell lines, in which two *VSG* genes are constitutively expressed in tandem from the same expression site. VSG double expressors were first generated in 1996 (Muñoz-Jordán et al., 1996), but the leveling of *VSG* mRNAs long remained undetected or anecdotal. We propose that trypanosomes have evolved an effective way of protecting the ER of becoming clogged by the dominant Pol I-transcribed *VSG* mRNA, by balancing the total amount of *VSG* mRNA to levels that support growth, even in the case that a VSG switch is unsuccessful (Batram et al., 2014; Zimmermann et al., 2017). We have shown that 50% of the wild type levels of *VSG* mRNA is sufficient for VSG coat formation and hence, the parasites seem to operate on the safe side. In the present study, we have explored some basic features of the *VSG* mRNA that are involved in mRNA balancing. We first asked if the process requires a *T. brucei* VSG and hence if it is species-specific. The related African trypanosomes *T. congolense* and *T. vivax* also express VSGs on their cell surface and the VSG coat is subject to antigenic variation, however, the VSG proteins may be divergent from *T. brucei*. While we have studied VSGs from both species, for the present study, we decided to challenge *T. brucei* with overexpression of two *T. vivax* VSGs. This turned out to be more complex than we thought.

Inducing the expression of the *T. vivax ILDat1*.*2 VSG*, resulted in a rapid reduction of the endogenous *T. brucei VSG221* mRNA amounts, similar to the results obtained when *T. brucei* VSGs were overexpressed (Batram et al., 2014). However, inducing the expression of another *T. vivax VSG, ILDat2*.*1*, did not affect the endogenous *VSG221* mRNA levels, despite the efficient production of *ILDat2*.*1* transcript. We initially thought that this result might reflect the high phenotypic flexibility of trypanosomes. However, we had never observed high level expression of *VSG* mRNA not resulting in mRNA balancing. Therefore, we surmised that the *ILDat2*.*1* transcript lacked an essential feature. The first candidate was the ER signal sequence, as we found that the *ILDat2*.*1* transcript was not translated. Whether the BSF trypanosomes rely on the SRP-dependent or independent pathways for translocation of VSGs has not been experimentally investigated (Manna et al., 2014). It has been suggested, however, that procyclic trypanosomes and *S. cerevisiae* appear to rely on the SRP-independent pathway for the translocation of GPI-anchored secretory proteins, and the SRP-dependent pathway for transmembrane-bearing proteins (Ast et al., 2013; Goldshmidt et al., 2008). In *S. cerevisiae*, cytosolic factors including the Heat shock protein 40 (Hsp40), yeast dnaJ protein 1 (Ydj1), and Hsp70 are all involved in the SRP-independent translocation of GPI-anchored proteins (Ast et al., 2013). The presence of Ydj1 homologs in *T. brucei* indicates functional similarity, and therefore these factors could be involved in regulating *VSG* mRNA translocation.

Indeed, the expression of transcript of a hybrid ILDat2.1 *VSG* reporter containing a bona fide *T. brucei* VSG ER import signal resulted in the expected balancing of endogenous *VSG221* mRNA, proving that the process can be triggered by expression of a non-*T. brucei* VSG. The second insight from this experiment was that balancing is most probably independent from translation, because, although the ER signal was functional, still ILDat2.1 protein appeared to be poorly made. Lastly, the experiment strongly suggested that for mRNA balancing to operate, the transcripts need to be targeted to the ER.

The next questions we asked were (i) if the ER-signal must be VSG-specific and (ii) if mRNA balancing requires a VSG at all. The ER import signal peptides of GPI-anchored *T. brucei* proteins VSG, BARP, EP1 and FHR all conform to the general organization of signal peptides (Heijne, 1998; Walter & Johnson, 1994). In fact, we found that the ER import signals of BARP, EP1 and FHR can target VSGs to the ER. Interestingly, the *VSG* mRNA was differentially regulated in VSG double expressor cell lines expressing ectopic *VSG121* with BARP or FHR ER import signals, as compared to the cells expressing ectopic *VSG121* with its native or EP1 ER import signal; the mRNA of the endogenous *VSG221* and ectopic *VSG121* with its native or EP1 ER import signal were expressed on a 70:30 basis (Supplementary Figure 2) whereas an equal expression at the transcript level was observed in the cell lines expressing ectopic *VSG121* with BARP and FHR ER import signals in addition to the endogenous *VSG221*. These results agree with previous findings which showed that ER import signals can varyingly influence the translocation and processing of substrates (Levine et al., 2005). It has been reported that specific motifs in the h-region can determine the efficiency of translocation of substrates into the ER of *T. brucei* (Duffy et al., 2010). Therefore, possibly the variations in *VSG* mRNA balancing were due to differences in the h-region motifs of the reporter VSG ER signals. Importantly, however, the experiments clearly showed that the VSG ER signal does not harbor special features required for the balancing of *VSG* mRNA.

To address the question whether the “signal” for mRNA balancing resides in the VSG open reading frame, we used ER-targeted and cytosolic GFP reporters coupled to the *VSG* 3′UTR to ensure high levels of expression. In agreement with all data obtained so far, we showed that high expression levels of ER-targeted *GFP* from the rDNA spacer locus resulted in a *trans*-regulation of the *VSG* mRNA. This regulation was absent when the *GFP* reporter was not targeted to the ER. The GFP reporter expression experiments showed that *VSG* mRNA balancing is neither dependent on the VSG open reading frame nor on the VSG ER import signal. This suggests that the expression of any highly abundant ER-targeted mRNA is sufficient for balancing the *VSG* mRNA amounts.

The only common denominator in all our experiments was the presence of an enigmatic 16-mer motif in the 3′UTR of the transcripts. This motif is essential for *VSG* mRNA stability in bloodstream stage parasites and is 100% conserved in all *T. brucei VSG* mRNAs. Interestingly, it is completely absent from the genomes of the related African trypanosome species *T. congolense* and *T. vivax*, which also have a VSG surface coat. The F-box RNA binding protein (CFB2) has been suggested as a limiting factor which interacts with the 16-mer motif in the VSG 3′UTR, thereby actively regulating *VSG* mRNA abundance (do Nascimento et al., 2020). As an alternative mechanism, *VSG* mRNA levels might be modulated by a feedback that is dependent on the production of functional GPI-anchored VSG protein (Maudlin et al., 2021).

In our experiments, the ectopic *VSG121* and *ILDat2*.*1 VSG* reporters containing the native *ILDat2*.*1* VSG ER import signal, which does not appear to function in *T. brucei*, were fused to complete *T. brucei* VSG 3′UTRs with intact 16-mer motifs. High levels of 16-mer-containing mRNA were expressed in the cytosol. However, levels of endogenous *VSG* mRNA were not downregulated, suggesting that the total *VSG* mRNA greatly exceeded the wild type amounts in these cells. This agrees with the study by Maudlin et al., which shows that *VSG* mRNA can be expressed above the wild type levels (Maudlin et al., 2021). These observations also imply that, in principle, the available CFB2 protein pool is sufficient to interact with the 16-mer of a second highly expressed *VSG* and can thus stabilize the mRNA. This points against a simple counting mechanism. We, therefore, propose that though the interaction between the 16-mer and CFB2 is essential for high expression and stability of *VSG* mRNA (do Nascimento et al., 2020; Ridewood et al., 2017), the steady-state *VSG* mRNA amount is regulated at a different level.

By introducing a PTC just before the ectopic VSG221 GPI signal, Maudlin et al., suggested that regulation of the *VSG* mRNA required GPI anchoring of the ectopic VSG (Maudlin et al., 2021). However, this data is inconsistent with ours as it implies that the modulation of *VSG* mRNA is not at the level of ER targeting, but rather downstream. Our data clearly show that an ER-targeted *VSG* transcript with a defective GPI-signal will trigger the same mRNA balancing as a fully functional mRNA. Previously published studies reported a 50:50 expression of *VSG* mRNA when the wild type *VSG* was integrated into the VSG221 ES (Muñoz-Jordán et al., 1996; Smith et al., 2009), whereas Maudlin et al., express the second VSG from the VSG121 ES and the VSG221:VSG121 levels have a 1:3 ratio (Maudlin et al., 2021). Additionally, expressing ectopic VSG221 with PTCs at different locations activated a pathway that increased the total *VSG* mRNA amounts. These variations in the modulation of *VSG* mRNA levels when different ES are used suggests there might be additional intricacies for the regulation of *VSG* mRNA.

In summary, we have shown that *VSG* mRNA balancing is independent of VSGs and of translation, but clearly dependent on high levels of ER-targeted transcripts. In view of the biology of trypanosomes this makes sense. Trypanosomes lack transcriptional control and highly abundant surface proteins are expressed by Pol I (Clayton, 2002; Günzl et al., 2003). During antigenic variation, mRNA balancing would cause an exchange of the *VSG* mRNA population before the completion of an expression site switch. During developmental progression to the procyclic insect stage, high level expression of ER-targeted procyclin mRNA would balance the amount of *VSG* mRNA present in the cell. This would likewise be true when procyclin is replaced with BARP in the next step of life cycle progression. Thus, trypanosomes exploit a simple but very effective system of posttranscriptional transcript balancing that allows for high phenotypic plasticity and robustness. In some aspects, transcript balancing in trypanosomes is reminiscent of a phenomenon called transcript buffering, which was first described in yeast (Haimovich et al., 2013; Sun et al., 2012). Transcript buffering involves a complex interplay that regulates transcription rates in the nucleus and mRNA decay in the cytoplasm to ensure steady-state mRNA levels are maintained in the cell pe (Pérez-Ortín et al., 2012; Shalem et al., 2008; Sun et al., 2013). This implies that there is a crosstalk between transcription and mRNA decay. Transcript buffering has further been demonstrated in mammalian cells (Abernathy et al., 2015; Singh et al., 2019), indicating a conserved process in eukaryotes. Transcript buffering can either be gene-specific or global and is possibly regulated by distinct mechanisms. How signal for crosstalk between transcription and mRNA decay is perceived, and its directionality is not well understood. In yeast, central players in transcript buffering include transcription initiation and the mRNA decay pathway factors (Timmers & Tora, 2018). In trypanosomes, the situation is certainly different from yeast and mammals, as transcription is not regulated. However, the basic need for transcript buffering also applies. We call the phenomenon transcript balancing as just two players are involved; in bloodstream stage trypanosomes these are two populations of *VSG* mRNAs during antigenic variation. The intriguing simplicity of the trypanosome system opens avenues that the opisthokont models lack. It has been suggested that mRNAs with longer half-lives are ideal for transcript buffering studies (Timmers & Tora, 2018). Therefore, the high-level expression and the long half-life of *VSG* mRNA can be harnessed to study transcript buffering in the tractable *T. brucei* system.

## Methodology

### Cultivation and genetic manipulation of trypanosomes

All cell lines generated in this study are based on the monomorphic *T. brucei* 427 MITat1.2 13-90 bloodstream-form parasites expressing the T7-polymerase and tetracycline repressor (Wirtz et al., 1999). The cells were maintained below 1 × 10^6^ cells/ml in HMI-9 medium supplemented with 10% heat-inactivated fetal calf serum (Sigma-Aldrich) at 37 °C and 5% CO_2_. For transfections, 10 µg of linearized DNA was transfected into 3.0 × 10^7^ mid-log phase cells in Amaxa Basic Parasite Nucleofector Solution 1 using the X-001 program of an Amaxa Nucleofector II (Lonza, Switzerland). Hygromycin, G418, blasticidin and phleomycin were used at 5, 2.5, 5, and 1 µg/ml, respectively, for the selection of recombinant cell lines.

Monomorphic wild type *T. brucei* Lister 427 cells expressing VSG121 was cultivated in HMI-9 medium supplemented with 10% heat-inactivated fetal bovine serum (Sigma-Aldrich) at 37 °C and 5% CO_2_ without antibiotics.

### Plasmid construction and generation of cell lines

The GFP coding sequence was amplified from plasmid p3822 (kindly provided by M Carrington, Cambridge, UK) using primers CFP_U.HindIII and 04_GFP_BamHI_L and ligated into pBSK II (+) plasmid with SmaI and BamHI restriction sites. The VSG 3’UTR was amplified from plasmid pBSK.M1.6-198 with primers M1.63’UTRBglII_U and M1.6_full_L and ligated after the GFP coding sequence using Bglll and BamHI restriction sites. The *GFP:VSG121 3’UTR* hybrid was excised using HindIII and XbaI and cloned into pLew82v4 (24,009; Addgene plasmid) to generate the pLew.GFP-198 construct. This construct was then linearized with NotI and transfected into the 13-90 parental cell line to generate the 221^ES^.GFP^tet^ cell line. For the addition of an ER-import signal to the *GFP* reporter, the *EP1* ER import signal sequence (EP^SP^) was excised from plasmid pLew.SP:GFP:GPI-198 with BstEII. The EP^SP^ was ligated into BstEII linearized pLew.GFP-198 plasmid resulting in the pLew.EP^SP^:GFP-198 plasmid which was NotI-linearized and transfected into 13-90 cells to generate the 221^ES^.EP^SP^:GFP^+tet^ cell line.

*ILDat1*.*2* and *ILDat2*.*1* coding sequence fused to *MITat1*.*1 VSG* 3’UTR were synthesized with EcoRI restriction sites at the 5’ and 3’ends and cloned into pBSK II (+) resulting in plasmid pBSK.ILDat1.2 and pBSK.ILDat2.1, respectively. The inserts were excised using HindIII and SmaI, followed by ligation into pLew82v4 plasmid that was linearized sequentially with XhoI (followed by refilling the overhangs with the Klenow fragment) and HindIII. The pLew.ILDat1.2 and pLew.ILDat2.1 plasmids were NotI-linearized and transfected into *T. brucei* 13-90 cells to generate the 221^ES^.ILDat1.2^tet^ and 221^ES^.ILDat2.1^tet^ cell lines, respectively.

Chimeric sequences were generated by PCR-driven overlap extension (Heckman and Pease, 2007). To replace the native *ILDat2*.*1* ER import signal with a *VSG121* ER import signal, *ILDat2*.*1:M1*.*1 3’UTR* sequence without the ER import signal and the *VSG121* ER import signal were amplified from plasmid pBSK.ILDat2.1 and pLew.121wt, respectively, using long primers that covered the overlap region of the two sequences. Products of the two PCR reactions were used as the template in a subsequent PCR to fuse the two fragments using primers annealing to the start of *VSG121* ER import signal and the end of *ILDat2*.*1:M1*.*1 3’UTR* sequence. The fused fragment was subcloned into pJet1.2 and pBSK II (+). The insert was excised and ligated into the pLew82v4 vector as described above to generate the 221^ES^.121^SP^:ILDat1.2^tet^ cell line. A similar approach was used to replace the native *VSG121* ER import signal with that of *ILDat2*.*1 VSG* from plasmid pJET.121 and pBSK.ILDat2.1, respectively. Transfection of the linearized construct into the parental cells generated the 221^ES^.ILDat1.2^SP^:121^tet^ cell line.

For constitutive expression of the GFP reporter from the ES, the VSG121 3’UTR was amplified from plasmid pRS.121 (Batram et al., 2014) with primers flanked by PacI and BamHI overhangs. The amplified fragment was cloned into pJet1.2 and then excised with PacI and BamHI followed by ligation into p3845 (kindly provided by M Carrington, Cambridge, UK) linearized with the same enzymes to generate plasmid p3845.VSG121 3’UTR. The *GFP* ORF coupled to *VSG121* 3’UTR was amplified from the plasmid p3845.VSG121 using primers flanked with BamHI and HindIII restriction sites. The amplicon was cloned into pJET1.2 resulting in plasmid pJET1.2:Blas:GFP:UTR. The Blas:GFP:UTR fragment was excised with BamHI and HindIII and ligated into a modified pbRN6 plasmid (Janzen et al., 2004) linearized with the same enzymes yielding the plasmid pbRN6.GFP:UTR. The upstream integration region of the resultant plasmid was modified by extending the *VSG221* 3’UTR portion to include the native polypyrimidine tract. A 452 bp fragment of part of the *VSG221* ORF and 3’ UTR was amplified from pBSK.VSG221wt-full 3’UTR plasmid. The amplicon was cloned into pJET1.2, excised with SacI and BamHI, and ligated into pbRN6.GFP:UTR. The construct was linearized with SacI and SalI and transfected into the 13-90 cells to generate the 221^ES^.GFP cell line. Next, a *VSG121* ER-targeting signal was fused to the *GFP:UTR* sequence by sequential PCR from plasmid pbRN6.GFP:UTR using long primers as described above. The amplicon was cloned into pJET1.2 and excised using HindIII and Eco1051 for cloning into HindIII and Eco1051-digested plasmid pbRN6.GFP:UTR. The generated pbRN6.121^SP^:GFP:UTR construct was transfected into 13-90 cells to generate the 221^ES^.121^SP^:GFP cell line.

For constitutive expression of VSG 121 downstream of the expression site resident VSG 221, the *VSG121* ORF was amplified from plasmid pRS.121 by PCR. The PCR product was blunted and ligated into pJET1.2:Blas:GFP:UTR after removal of the *GFP* ORF with PacI and PaeI restriction enzymes. Next, the Blas:VSG121:UTR was transferred from the pJET1.2:Blas:VSG121:UTR into a modified pbRN6 and the upstream integration region extended as described above to generate pbRN6.M1.6wt plasmid. The construct was linearized with SacI and SalI and transfected into the 13-90 cells. To replace the *VSG121* ER import signal with FHR signal, the region upstream of the *VSG121* gene and the *VSG121* sequence minus the signal peptide was amplified from plasmid pbRN6.M1.6wt using primers that contained the full signal sequence and fused as described above. Replacement of the *VSG121* ER import signal with the BARP and EP1 signal sequences was done in a similar manner. The reporter sequences were cloned into pJET1.2, excised using EcoRI and HindIII, and cloned into pbRN6.M1.6wt plasmid that was linearized in the same way. Transfection of SacI and SalI linearized constructs into 13-90 cells created the 221^ES^.FHR^SP^:121 and the 221^ES^.BARP^SP^:121 cell lines. Unless otherwise stated, all enzymes were obtained from Thermo Fisher Scientific (USA). The primers are available on request.

## RNA analysis

Total RNA was extracted from 1 × 10^8^ parasites using the RNeasy Mini Kit (Qiagen, Netherlands) as per the manufacturer’s instructions. For quantification of mRNA, 3 µg of total RNA was denatured with glyoxal at 50 °C for 40 min and transferred onto N-Hybond nitrocellulose membrane (GE Healthcare, UK) using a Manifold Dot blotter (Schleicher & Schuell, Germany). The blots were hybridized overnight at 42 °C with oligonucleotide probes labelled with fluorescent IRDye 682 (*VSG121*-probe: GGCTGCGGTTACGTAGGTGTCGATGTCGAGATTAAG; *VSG221*: CAGCGTAAACAACGCACCCTTCGGTTGGTCGTCTAG; *GFP*: GCCGTTCTTCTGCTTGTCGGCCATGATATAGA; *ILDat1*.*2 and 2*.*1*: TAGGATATCAAGCTTGTGAATTTTACTTTTTGG) or IRDye 782 (*β-tubulin:* ATCAAAGTACACATTGATGCGCTCCAGCTGCAGGTC). Imaging and quantification of fluorescence was done using the Li-Cor Odyssey or Li-Cor Odyssey CLx system (Li-Cor, Netherlands) and Image Studio Lite.

### Protein analyses

Trypanosomes were resuspended and lysed in protein sample buffer to yield equivalents of 2 × 10^5^ cells/µl. For quantification of VSG proteins by SDS-PAGE or Western blots, 5 µl of the protein sample was resolved on 12.5% SDS-PAGE gels and transferred onto nitrocellulose membrane. For quantification by dot blots, 3 µl of the protein sample was blotted on the nitrocellulose membrane. The membranes were blocked for 1 h at room temperature with 5% (w/v) milk powder in PBS. Next, the membranes were incubated in polyclonal primary antibodies diluted in PBS containing 1% (w/v) milk powder and 0.1% (v/v) Tween 20: rabbit anti-VSG221 1:5,000; rabbit anti-VSG121 1:2,000 and mouse monoclonal anti-PFR (L13D6) 1:20. After wash steps, the membranes were incubated in IRDye 782 labelled goat-anti-rabbit and IRDye 682 labelled goat-anti-mouse secondary antibodies diluted in PBS containing 1% (w/v) milk powder and 0.1% (v/v) Tween 20: 1:10,000. Imaging and quantification were done with the Li-Cor Odyssey system.

### Fluorescence microscopy

Parasites were fixed in a final concentration of 4% w/v formaldehyde and 0.05% v/v glutaraldehyde overnight at 4 °C. After fixation, the cells were washed twice with PBS and stained with 1 µg/ml of DAPI immediately before imaging. The cells were imaged using an iMIC widefield fluorescence microscope (FEI Photonics, Germany) fitted with a CCD camera (Sensicam qe, pixel size 6.45 μm, PCO, Germany) using a 100x (NA 1.4) objective (Olympus, Germany) and the filter cubes ET-GFP and DAPI (Chroma Technology CORP, USA). The set up was controlled by the Live acquisition software (FEI Photonics, Germany). Alternatively, trypanosomes were viewed with an automated DMI6000B wide field fluorescence microscope (Leica Microsystems, Germany) equipped with a DFC365FX camera (pixel size 6.45 µm) and a 100x oil objective (NA 1.4). The images are displayed as maximum intensity projections of Z-stacks. Image analysis was carried out using ImageJ (National Institute of Health).

## Acknowledgements

We thank Susanne Kramer for advice and critical reading of the manuscript. EOA is funded by the German Academic Exchange Service (DAAD). ME is supported by DFG grants EN305, SPP1726 (Microswimmers – From Single Particle Motion to Collective Behaviour), GIF grant I-473-416.13/2018 (Effect of extracellular *Trypanosoma brucei* vesicles on collective and social parasite motility and development in the tsetse fly), GRK2157 (3D Tissue Models to Study Microbial Infections by Obligate Human Pathogens) and the BMBF NUM Organo-Strat. ME is a member of the Wilhelm Conrad Roentgen Center for Complex Material Systems (RCCM).

## Supplementary Material

**Figure 1:**
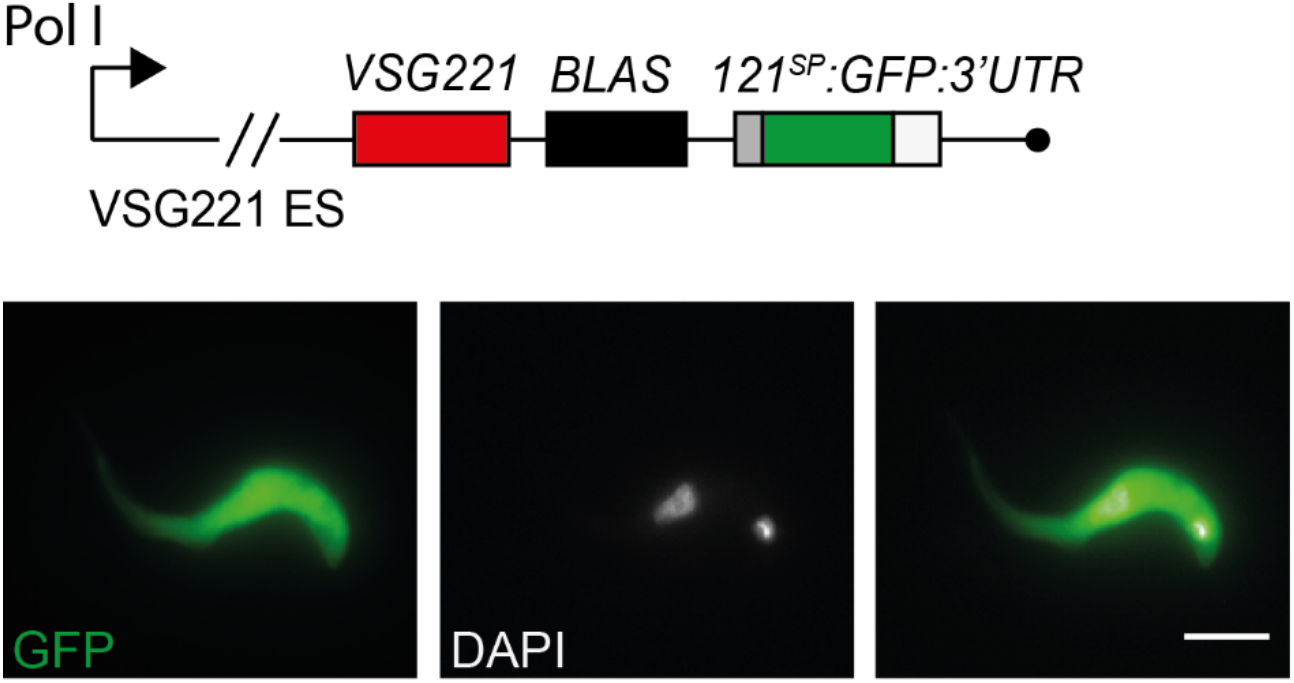
Expression of ER-targeted GFP reporter downstream of the active ES resident VSG221. Schematic of the *GFP* reporter construct used to generate the 221^ES^.121^SP^:GFP cell line and fluorescence imaging of fixed cells constitutively expressing GFP from the 221 ES. GFP fluorescence is shown in green while the nucleus and kinetoplast were stained with DAPI (white). Scale bar: 5 µm.

**Figure 2:**
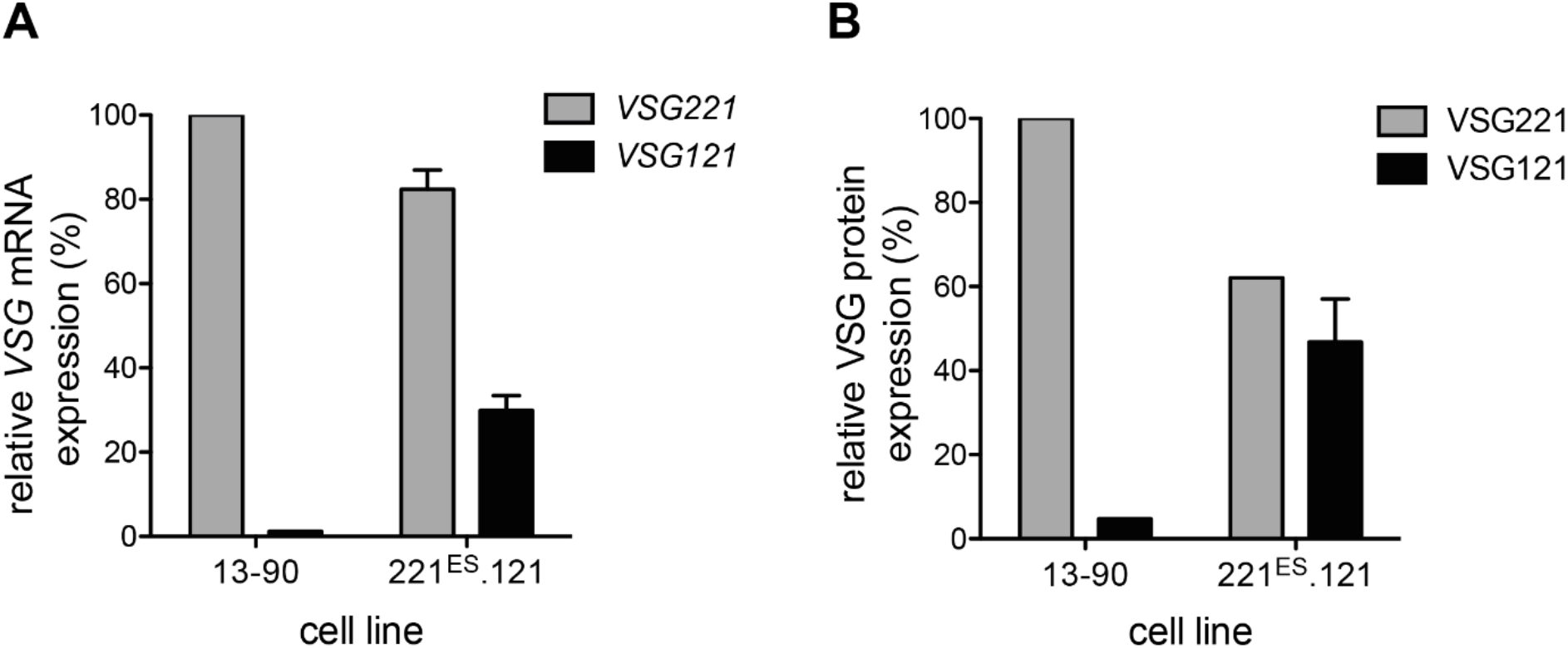
Expression of VSG121 downstream of the active BES-resident VSG221. (A) Quantification of *VSG221* and *VSG121* mRNA and (B) protein expression in the 221ES.121 cell line. *VSG* mRNA and protein were normalized to *β-tubulin* and PFR protein, respectively. Expression levels are presented as percentages relative to VSG expression in the parental 13-90 cells and wild type VSG121 for VSG221 and VSG121, respectively. Values are given as means and the error bars shows ± SD of three independent clones.

**Figure 3:**
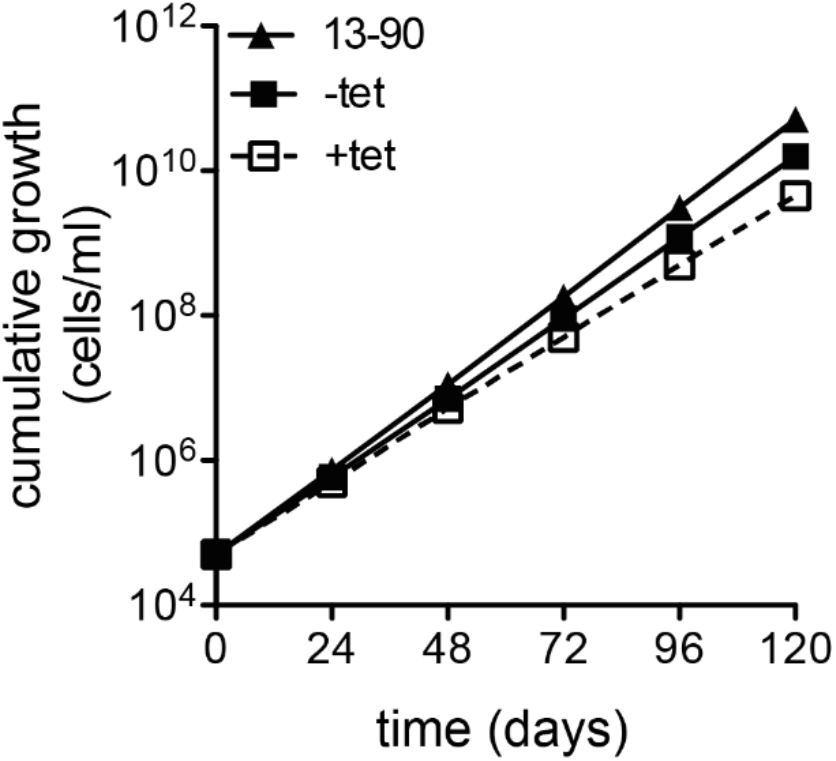
Cumulative growth curve of the cell line expressing ILDat1.2 VSG whereby the native GPI signal was replaced with one from a *T. brucei* VSG. The growth was analyzed for five days in the presence and absence of tetracycline. The parental 13-90 cells served as the control. Three independent clones were analyzed, and the error bars represent standard deviation of the means.

**Table 1.**
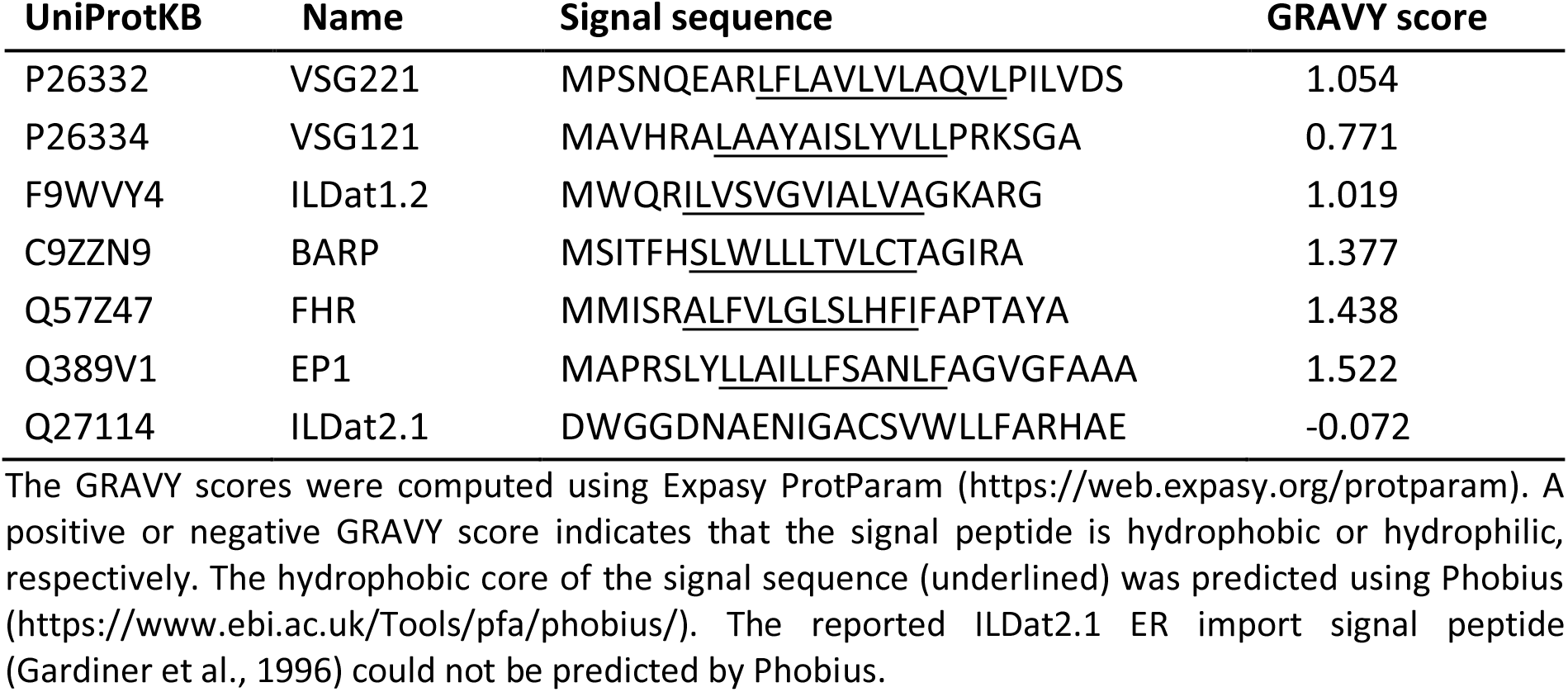
The grand average hydropathy (GRAVY) scores of select ER import signals. The reported ILDat2.1 ER import signal is hydrophilic.

